# Direct activation of endothelial cells by SARS-CoV-2 nucleocapsid protein is blocked by Simvastatin

**DOI:** 10.1101/2021.02.14.431174

**Authors:** Yisong Qian, Tianhua Lei, Parth S. Patel, Chi H Lee, Paula Monaghan-Nichols, Hong-Bo Xin, Jianming Qiu, Mingui Fu

## Abstract

Emerging evidence suggests that endothelial activation plays a central role in the pathogenesis of acute respiratory distress syndrome (ARDS) and multi-organ failure in patients with COVID-19. However, the molecular mechanisms underlying endothelial activation in COVID-19 patients remain unclear. In this study, the SARS-CoV-2 viral proteins that potently activate human endothelial cells were screened to elucidate the molecular mechanisms involved with endothelial activation. It was found that nucleocapsid protein (NP) of SARS-CoV-2 significantly activated human endothelial cells through TLR2/NF-κB and MAPK signaling pathways. Moreover, by screening a natural microbial compound library containing 154 natural compounds, simvastatin was identified as a potent inhibitor of NP-induced endothelial activation. Remarkablely, though the protein sequences of N proteins from coronaviruses are highly conserved, only NP from SARS-CoV-2 induced endothelial activation. The NPs from other coronaviruses such as SARS-CoV, MERS-CoV, HUB1-CoV and influenza virus H1N1 did not affect endothelial activation. These findings are well consistent with the results from clinical investigations showing broad endotheliitis and organ injury in severe COVID-19 patients. In conclusion, the study provides insights on SARS-CoV-2-induced vasculopathy and coagulopathy, and suggests that simvastatin, an FDA-approved lipid-lowering drug, may benefit to prevent the pathogenesis and improve the outcome of COVID-19 patients.

## INTRODUCTION

The emergence of severe acute respiratory syndrome coronavirus 2 (SARS-CoV-2), which causes the coronavirus disease 2019 (COVID-19), triggered a global pandemic that has led to an unprecedented worldwide public health crisis^1^. Before SARS-CoV-2, two other highly pathogenic coronaviruses emerged in the past two decades, including severe acute respiratory syndrome coronavirus (SARS-CoV)^2^ and Middle East respiratory syndrome coronavirus (MERS-CoV)^3^. In addition, four endemic human coronaviruses (i.e., OC43, 229E, NL63 and HKU1) cause common cold respiratory diseases^4^. COVID-19 is characterized by progressive respiratory failure resulting from diffuse alveolar damage, inflammatory infiltrates, endotheliitis, and pulmonary and systemic coagulopathy forming obstructive microthrombi with multiorgan dysfunction^5-8^. Pathological findings of cell swelling, severe endothelial injury, disruption of intercellular junctions, and basal membrane contact loss in COVID-19 patients imply that the destruction of endothelial cells (ECs) leads to pulmonary vascular endotheliitis and alveolar capillary microthrombi^9-12^. Together, emerging evidence suggests that endothelial activation is an early hallmark of multiorgan damage in patients with COVID-19^13^. Moreover, thrombotic complications are a relevant cause of death in patients with COVID-19^14^. Therefore, understanding the molecular mechanisms of the endothelial activation caused by SARS-CoV-2 and pathways involved in the regulation of endothelial dysfunction could lead to new therapeutic strategies against COVID-19.

SARS-CoV-2 infects the host using angiotensin converting enzyme 2 (ACE2) as its receptor^15^. ACE2 is an integral membrane protein that is expressed by airways and lung alveolar epithelial cells, enterocytes, and vascular endothelial cells^16^. Though the primary target tissues of SARS-CoV-2 are airways and lungs, there is also evidence of direct viral infection of endothelial cells and diffuse endothelial inflammation in COVID-19 disease^17^. Moreover, vulnerable patients with pre-existing endothelial dysfunction, which is associated with male sex, smoking, hypertension, diabetes and obesity and established cardiovascular disease, are associated with adverse outcomes in COVID-19^18^. Together, these clinical findings suggest that endothelial cells play a central role in the pathogenesis of ARDS and multi-organ failure in patients with COVID-19. Therefore, the vascular system is increasingly being addressed as a major therapeutic target for defeating COVID-19.

The potential molecular mechanisms that SARS-CoV-2 induces endothelial activation, dysfunction and injury may contribute to the direct toxic effect of viral proteins. The genome of SARS-CoV-2 encodes 29 viral proteins including 16 non-structure proteins (NSP1-NSP16), 4 structure proteins including spike (S), membrane (M), nucleocapsid (N) and envelope (E) proteins and 9 accessory proteins^19, 20^. Though their functions in viral lifecycle are increasingly studied, their impacts on host cells are largely unknown. To determine whether distinct viral proteins can induce endothelial activation, recombinant SARS-CoV-2 proteins were evaluated for their potential to activate human endothelial cells. We found that N protein potently induced endothelial activation via Toll-like receptor 2 (TLR2)/NF-κB and MAPK signal pathways. To identify potential therapeutic agents targeting N protein-induced endothelial activation, a natural microbial compound library containing 154 natural compounds was screened and a single drug, simvastatin, was identified as specific inhibitor of N protein-induced endothelial activation. These results suggest that N protein released from SARS-CoV-2-infected cells may contribute to the broad activation of endothelium and tissue inflammation and simvastatin may benefit to prevent the viral infection-induced pathogenesis and improve the outcome of COVID-19 patients.

## RESULTS

### Identification of nucleocapsid protein (NP) as a potent inducer of human endothelial cell activation

To understand the molecular mechanisms that SARS-CoV-2 induces endothelial activation, we purchased eight recombinant SARS-CoV-2 viral proteins, including three structural proteins (S, N, and E proteins) and five non-structural proteins (NSP1, NSP3, NSP5, NSP7 and NSP8). The proteins were added to human primary lung microvascular endothelial cells (HLMECs) for 8 hours. Western blotting was used to analyze the expression of ICAM-1 and VCAM-1, the markers of endothelial cell activation. As shown in **Fig.1a**, N protein significantly induced the expression of ICAM-1 and VCAM-1. TNFα is the most potent endogenous inducer of endothelial activation, which serves as a positive control. To further confirm the effect of N protein on endothelial cell activation, we incubated HLMECs with different doses of N protein or for different incubation periods. As shown in **Fig.1b**, N protein induced ICAM-1 expression at 4 hours and continued to increase up to 24 hours, which was similar to the expression pattern induced by TNFα. Similar pattern of expression of VCAM-1 was also induced by N protein. However, the expression of VE-cadherin, an EC junction structure protein, was not changed. In addition, N protein significantly induced ICAM-1 expression at 0.05 μg/ml and more potent at 1 μg/ml, which is comparable with the effect of TNFα at 10 ng/ml (**Fig.1c**). Next, we tested whether N protein also induced activation of other endothelial cells. Human umbilical vein endothelial cells (HUVEC), human aortic endothelial cells (HAEC), human coronal artery endothelial cells (HCAEC), human dermal microvascular endothelial cells (HDMEC), HLMEC, 293T, A549 and mouse lung vascular endothelial cells (MEC) were incubated with 1 µg/ml of N protein for 8 hours. As shown in **Fig.1d**, N protein significantly induced activation of all human endothelial cells tested, but not induced expression of ICAM-1 and VCAM-1 in 293T, A549 and mouse endothelial cells. We also probed the same blot using anti-mouse ICAM-1 and VCAM-1 antibodies, respectively, and confirmed that N protein did not induced mouse EC activation. To further confirm whether N protein induce the expression of adhesion molecules and proinflammatory cytokines at transcriptional level, we examined the changes of ICAM-1, VCAM-1, E-selectin, TNFα, MCP-1 and IL-1β mRNA levels in NP-treated HLMECs. As shown in **Fig.1e**, N protein potently induced the expression of adhesion molecules, inflammatory cytokines and chemokines, which is similar to the function of TNFα, a potent endogenous inducer of endothelial activation. The expression of ICAM-1 and VCAM-1 on the surface of endothelial cells contributes to the leukocytes adherence on endothelial cells^21,22^. To examine if N protein also induces the monocyte adherence to activated endothelial cells, HLMECs were stimulated with N protein (1 μg/ml) or TNFα (10 ng/ml) for 8 hours and then the Zombie Red fluorescent-labelled human primary monocytes (Lonza) were co-cultured with the activated endothelial cells for 1 hour. After washed, the adherent cells were visualized by a fluorescent microscopy and counted in a double-blind way. As shown in **Fig.1f**, both N protein and TNFα significantly increased monocyte adherence on ECs, compared with that in control group. Taken together, these results suggest that N protein is a potent inducer of endothelial cell activation, and it may play a key role in SARS-CoV-2-induced lung inflammation and multi-organ failure.

**Figure 1.**
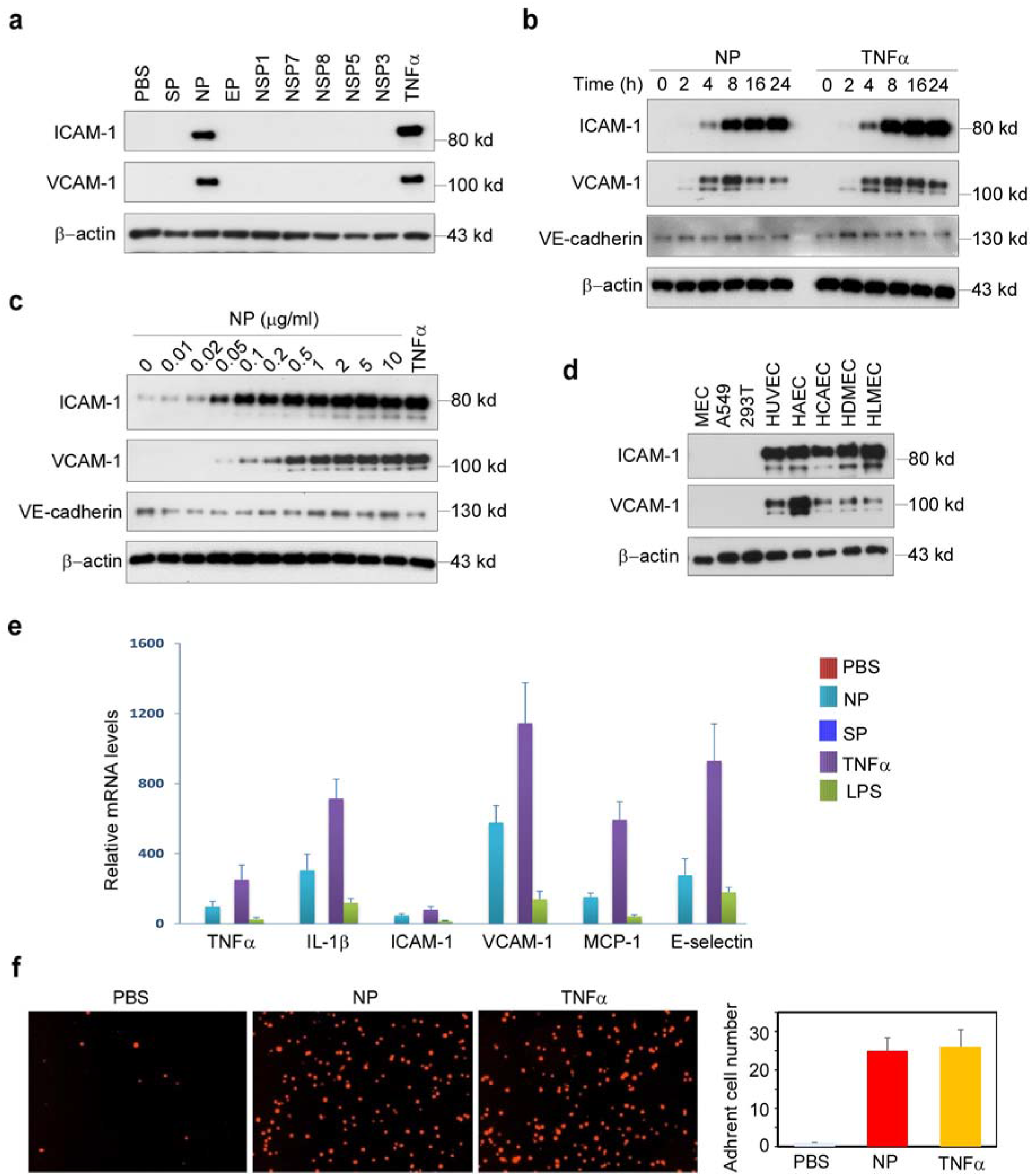
SARS-CoV2 nucleocapsid protein (NP) is a potent inducer of human endothelial cell activation. **(a)** HLMECs were incubated with SARS-CoV2 structural proteins (S, N, and E proteins, 1 μg/mL) and five non-structural proteins (NSP1, NSP3, NSP5, NSP7 and NSP8, 1 μg/mL) for 8 hrs. **(b)** HLMECs were treated with 1 μg/mL of NP or 10 ng/ml of TNFα for different incubation periods as indicated. **(c)** HLMECs were incubated with indicated concentrations of NP for 8 hrs. 10 ng/ml of TNFα serves as positive control. **(d)** Different cultured cells including mouse lung vascular endothelial cells (MEC), A549, 293T, HUVEC, HAEC, HCAEC, HDMEC and HLMEC were treated with NP (1 μg/mL) for 8 hrs. The expression of ICAM-1, VCAM-1 and VE-cadherin was detected by western blot. β-actin was served as loading control. **(e)** HLMECs were treated with PBS, NP (1 µg/ml), TNFα (10 ng/ml) or LPS (1 µg/ml) for 8 hours. The total RNA was isolated and QPCR was performed for measuring the mRNA levels of TNFα, ICAM-1, VCAM-1, MCP-1 and IL-6. **(f)** HLMECs were treated with PBS, NP (1 µg/ml) or TNFα (10 ng/ml) for 8 hours and co-cultured with Zombie Red-labeled THP-1 cells for 1h. After washing, the adherent cells were imaged and quantitatively analyzed.

### N protein activated NF-κB and MAPK signaling pathways in human endothelial cells

It is well known that the expression of ICAM-1 and VCAM-1 was controlled by MAPK and NF-κB signaling pathways^23,24^. To test if N protein can activate these signal pathways, HLMECs were incubated with NP (1 µg/ml) for 0, 15, 30, 45, 60 and 120 min, the cell lysates were analyzed by Western blot. As shown in **Fig.2**, NP treatment induced the phosphorylation of IKKs, p65 and IκBα, JNK, p38, and led to IκBα degradation. The time patterns of phosphorylation of these signaling molecules were similar to that of TNFα-induced activation of the signal pathways. Interested to note that S protein and LPS did not induce activation of NF-κB signaling, but weakly induced activation of MAPK pathways including ERK1/2, JNK and p38. These results suggest that N protein activated JNK, p38 and NF-κB signal pathways in human endothelial cells.

**Figure 2.**
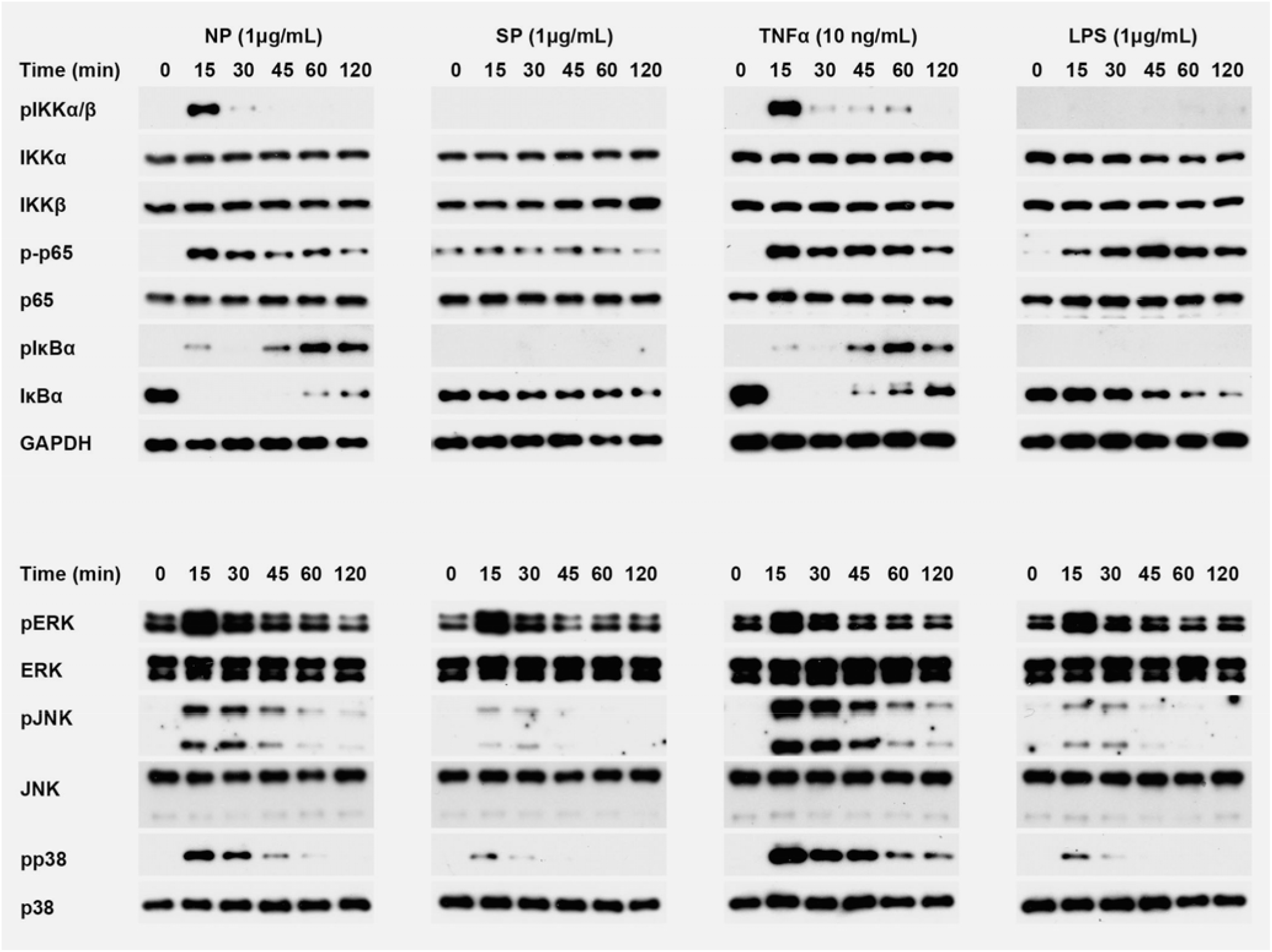
N protein activated NF-κB and MAPK signaling pathways in human endothelial cells. HLMECs were incubated with NP (1 µg/ml), SP (1 µg/ml), TNFα (10 ng/ml) and LPS (1 µg/ml) respectively for indicated times. The phosphorylation of IKKs, p65, IκBα, ERK, JNK and p38, as well as IκBα degradation were detected by western blot. GAPDH was served as loading control.

### N protein induced endothelial cell activation via TLR2-mediated signaling pathway

Next, we investigated how N protein activates NF-κB and MAPK signaling pathways in human endothelial cells. First we test if N protein acts on cell surface receptors or intercellular signal proteins. Pretreatment with the inhibitors of endocytosis such as Pitstop2 and Dynasore hydrate did not affect NP-induced expression of ICAM-1 and VCAM-1 (**Fig.3a**), suggesting that internalization of N protein is not required for its action on endothelial activation. Moreover, we transfected the expression plasmid encoding N protein into HLMECs. Though overexpression of N protein in endothelial cells, intercellular N protein did not induce the expression of ICAM-1 and VCAM-1 (**Fig.3b**). Furthermore, incubation of HLMECs with N protein for different times showed that N protein bound to endothelial cells in a time-dependent manner (**Fig.3c**). These results suggest that N protein may bind to a kind of receptors on endothelial cells and trigger the NF-κB and MAPK signal pathways. Next, HLMECs were pretreated with antagonists of TLR2 (CU-CPT22), TLR4 (LPS-RS) and IL-1R (IL-1R antagonist) for 1 hour and then incubated with 1 µg/ml of N protein for 8 hours. The expression of ICAM-1 and VCAM-1 was detected by Western blotting. As shown in **Fig.3a**, TLR2 antagonist (CU-CPT22) significantly blocked NP-induced expression of ICAM-1 and VCAM-1 in human endothelial cells, suggesting that N protein may bind to TLR2 to trigger the activation of NF-κB and MAPK and induce endothelial activation. **Fig.3d** further showed that CU-CPT22 dose-dependently inhibited NP-induced expression of ICAM-1 and VCAM-1. To further confirm that N protein is able to activate TLR2, both wild-type 293T cells (without TLR2 expression) and TLR2-overexpressed 293T cells were treated with or without N protein. As shown in **Fig.3e**, N protein did not induce phosphorylation of JNK and p38 in wild-type 293T cells. However, N protein significantly induced phosphorylation of JNK and p38 in 293T-TLR2 cells. Finally, as shown in **Fig**,**3a**, pretreatment with the inhibitors of IKK, JNK and p38 completely blocked NP-induced expression of ICAM-1 and VCAM-1. Taken together, these results suggest that N protein activates endothelial cells via TLR2-mediated NF-κB and MAPK signal pathways.

**Figure 3.**
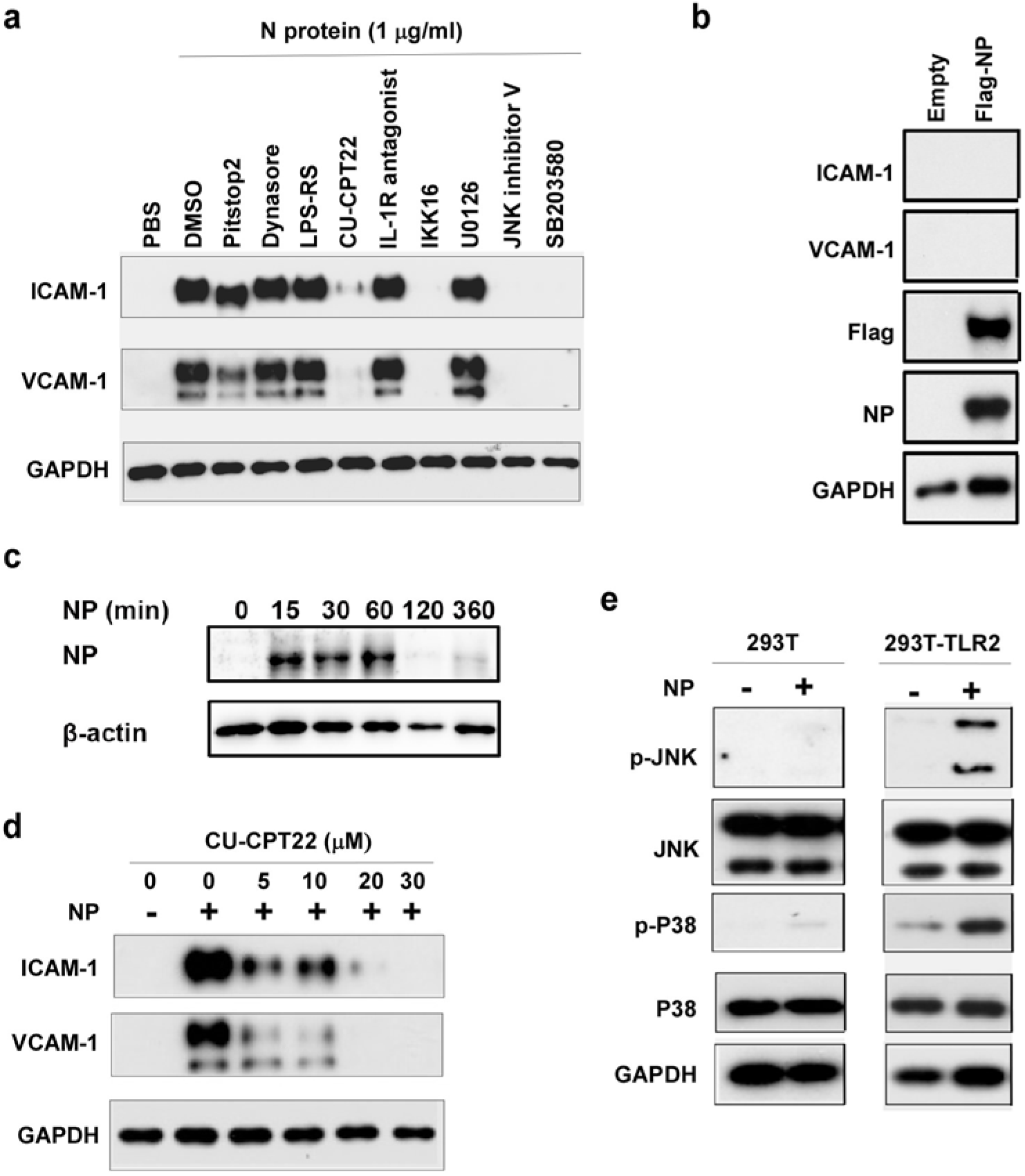
N protein induced endothelial cell activation via TLR2-mediated signaling pathway. **(a)** HLMECs were pretreated with inhibitors of endocytosis (Pitstop2, 12.5 μM and Dynasore hydrate, 12.5 μM) and antagonists of TLR4 (LPS-RS, 10 μg/mL), TLR2 (CU-CPT22, 20 μM), IL-1R (IL-1R antagonist, 20 μM), inhibitors of IKK (IKK16, 20 μM), ERK (U0126, 20 μM), JNK (JNK inhibitor V, 20 μM) and p38 (SB203580, 20 μM) for 1 h followed by treatment with NP (1 μg/mL) for 8 hrs. **(b)** The Flag control and Flag-NP expression plasmids were transfected into HLMECs by electroporation respectively. The whole cell lysate was harvested after 48 h transfection. The expression of ICAM-1, VCAM-1, NP and Flag was detected by western blot. **(c)** HMVECs were incubated with NP (1 μg/mL) for different time as indicated. After washing, the cells were harvested and NP was detected by western blot. **(d)** HLMECs were treated with indicated concentrations of CU-CPT22 for 1 h followed by treatment of NP (1 μg/mL) for 8 hrs. **(e)** Wild type (left) and TLR2-overexpressed 293T cells were treated with or without NP (1 μg/mL) for 15 min. The pJNK and pP38 were detected by western blot. GAPDH was served as loading control. All of experiments have been repeated at least one time.

### Identification of simvastatin as an effective inhibitor of N protein-induced endothelial activation

To screen the inhibitors of N protein-induced endothelial activation, 154 chemicals from microbial natural product library (Target Molecule, Wellesley Hills, MA) were added into individual wells of 48-well-plates at 30 μM 1 hour before the induction of N protein (1 μg/ml). The effect of chemicals on endothelial activation was measured by Western blot with anti-ICAM-1 antibody. Among 154 chemicals, we found that 12 chemicals showed significant inhibition of N protein-induced ICAM-1 expression, which include Simvastatin, Lovastatin, Rapamycin, Cyclosporine A, Menadione, 1, 4-Naphthoquinone, L-Thyroxine, Thiostrepton, Monensin, Amphotericin B, Gramicidin and Abamectin (**Fig.4**). The later six are antibiotics which are toxic for cells and used in animals, not suitable for human use. Menadione and 1, 4-Naphthoquinone are vitamin K derivatives, which have anti-bacteria, anti-viral, anti-inflammation activities. Rapamycin and Cyclosporine A are immunosuppressant. Simvastatin and lovastatin are FDA-approved lowering blood lipid drugs with anti-inflammatory roles.

**Figure 4.**
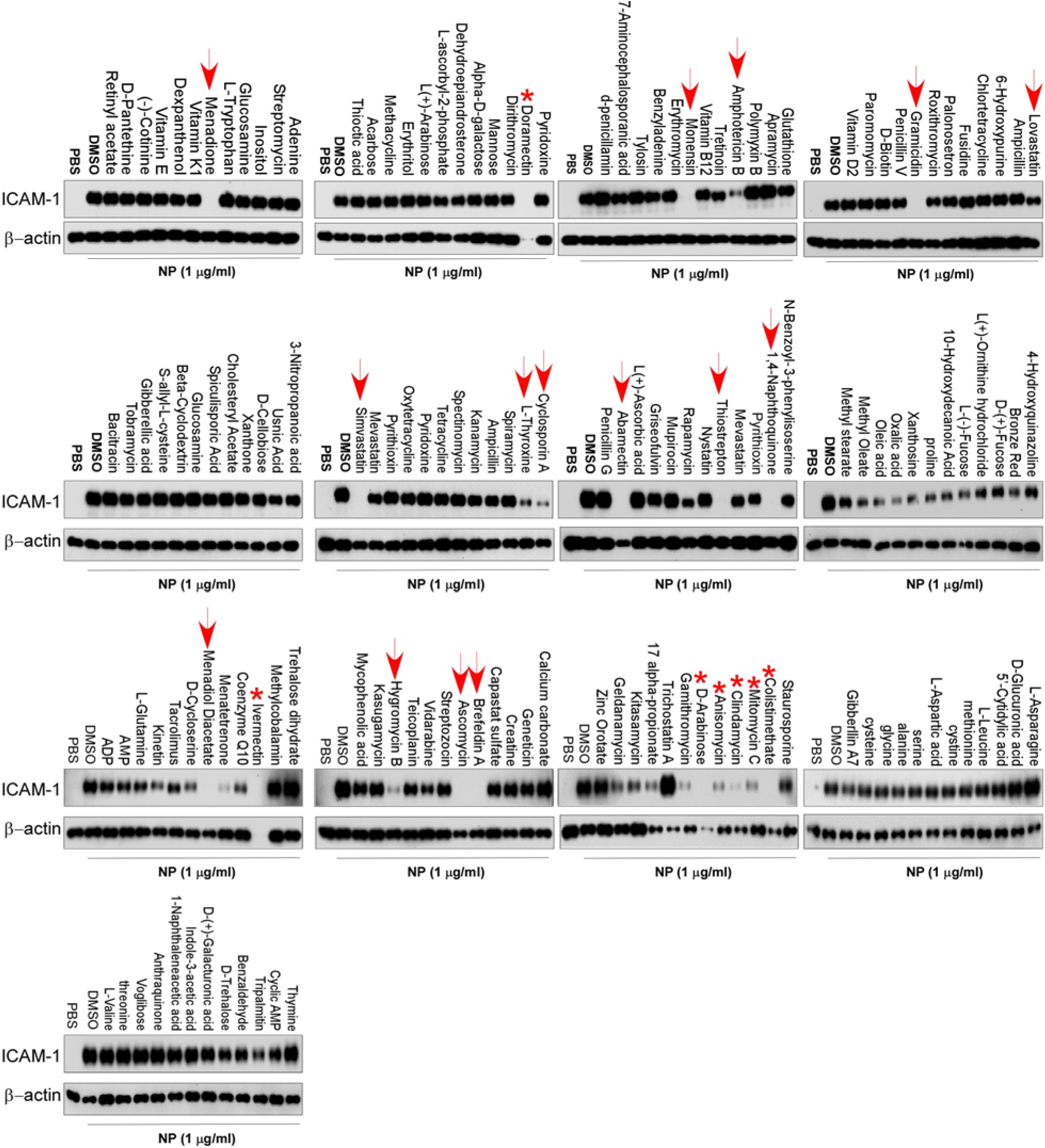
Screening of chemicals for inhibition of N protein-induced endothelial activation. A total of 155 chemicals from microbial natural product library were added into HLMECs at 30 μM 1 hour before the induction of N protein (1 μg/ml). The effect of chemicals on endothelial activation was measured by Western blot with anti-ICAM-1 antibody. β-actin was served as loading control. Arrow points out the effective compounds; stars point out the toxic compounds causing cell death. The experiments has been repeated for one more time.

### Simvastatin is an effective inhibitor of N protein-induced endothelial activation in vitro

To confirm the specific inhibitory role of simvastatin in N protein-induced endothelial activation, we compared the effect of simvastatin, lovastatin, atorvastatin, mevastatin and rosuvastatin. As shown in **Fig.5a**, among the five statins tested, simvastatin potently inhibited N protein-induced expression of ICAM-1 and VCAM-1. Consistent with the screening results, lovastatin showed mild effect on N protein-induced endothelial activation. The other three statins did not affect N protein-induced endothelial activation. We further confirmed the effect of simvastatin on N protein-induced endothelial activation in a dose-dependent manner **(Fig.5b**). Moreover, both simvastatin and lovastatin treatments significantly inhibited N protein-induced NF-κB activation (**Fig.5c**). Consistently, simvastatin pretreatment also blocked monocyte adhesion to the activated endothelial cells (**Fig.5d**). Simvastatin is a member of the class of hexahydronaphthalenes like lovastatin in which the 2-methylbutyrate ester moiety has been replaced by a 2,2-dimethylbutyrate ester group. Simvastatin is derived from lovastatin. Lovastatin is a fatty acid ester that is mevastatin carrying an additional methyl group on the carbobicyclic skeleton. The structures of simvastatin, lovastatin and mevastatin are very similar. Mevastatin does not have a methyl group, however, both simvastatin and lovastatin have this group, which suggest that the group may be important for its inhibitor effect on N protein-induced endothelial activation (**Fig.6**).

**Figure 5.**
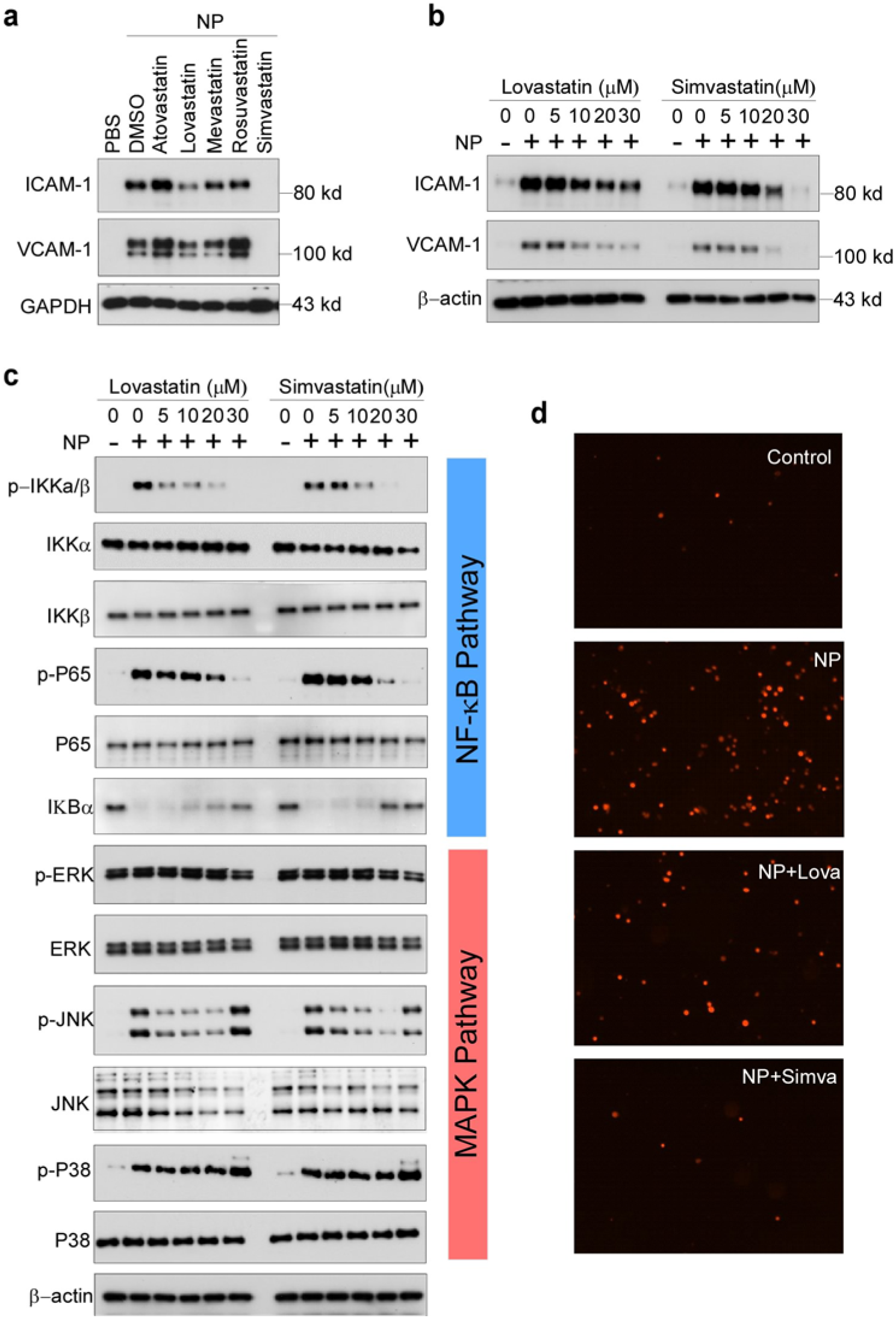
Simvastatin is an effective inhibitor of N protein-induced endothelial activation in vitro. **(a)** HLMECs were pretreated with simvastatin, lovastatin, atorvastatin, mevastatin and rosuvastatin at 30 μM for 1 h followed by treatment with NP (1 μ g/mL) for 8 hrs. **(b)** HLMECs were pretreated with indicated concentrations of Lovastatin and Simvastatin for 1 h followed by treatment with NP (1 μg/mL) for 8 hrs. The expression of ICAM-1 and VCAM-1 was detected by western blot. **(c)** HLMECs were pretreated with indicated concentrations of Lovastatin and Simvastatin for 1 h by treatment with NP (1 μg/mL) for 15 mins. The activation of NF-κB and MAPK signal pathways was detected by western blot. **(d)** HLMECs were pretreated with or without Lovastatin (Lova) or Simvastatin (Simva) followed by treatment with NP (1 µg/ml) for 8 hours and then co-cultured with Zombie Red-labeled THP-1 cells for 1 h. After washing, the adherent cells were imaged.

### The N protein from SARS-CoV-2 but not the other coronaviruses potently induced endothelial cell activation

There are seven types of coronaviruses infecting humans including SARS-CoV-2, SARS-CoV, MERS-CoV, OC43-CoV, HKU1-CoV, 229E-CoV and NL63-CoV. The previous three are highly pathogenic and cause severe problems for humans. However, the later four are less pathogenic and only cause common cold. Though SARS-CoV and MERS-CoV display high sequence similarity with SARS-CoV-2, the patients infected with SARS-CoV-2 are commonly affected by vascular injury and thrombosis formation^25^, which was not observed in the patients infected with SARS-CoV and MERS-CoV. To understand the molecular basis of the pathogenesis, we compared the effect of different N proteins from SARS-CoV-2, SARS-CoV, MERS-CoV, HKU1-CoV as well as H7N9 on endothelial activation. Interestingly, as shown in **Fig.7**, only the N protein from SARS-CoV-2 potently induced endothelial activation. The other N proteins, though their sequence is highly conserved, did not affect endothelial activation (**Fig.7**). These results may explain why SARS-CoV-2-infected patients developed severe vascular injury and thrombosis and affected many organs.

## DISCUSSION

Coronavirus disease 2019 (COVID-19), caused by SARS-CoV-2, is a worldwide challenge for health care system. The leading cause of mortality in patients with COVID-19 is ARDS and multi-organ failure. Emerging evidence suggests that pulmonary endothelial cells contribute to the initiation and progression of ARDS by altering vessel barrier integrity, promoting a pro-coagulation state, inducing vascular inflammation (endotheliitis) and mediating inflammatory cell infiltration. However, the molecular mechanisms underlying SARS-CoV-2-induced endothelial activation and vascular injury remain unclear. Though some reports indicated direct infection of endothelial cells by SARS-CoV-2 in human samples, emerging evidence suggests that ACE2 is not highly expressed in human endothelial cells and SARS-CoV-2 is not able to infect human endothelial cells in vitro^26^. Thus, the direct damage of endothelial cells by SARS-CoV-2 infection cannot explain the broad endothelial dysfunction in COVID-19 patients. The possible mechanisms by which SARS-CoV-2 infection causes endothelial activation may be attributed to the inflammatory and toxic roles of circulating viral proteins released from infected and lysed cells and inflammatory cytokines secreted from inflammatory and immune cells.

In the manuscript, we have screened the recombinant SARS-CoV-2 viral proteins that are able to activate human endothelial cells. We found that nucleocapsid protein (N protein) of SARS-CoV-2 potently activate endothelial cells through TLR2/NF-κB and MAPK signal pathways, by which N protein significantly induced the expression of ICAM-1 and VCAM-1 as well as other inflammatory cytokines and chemokines such as TNFα, IL-1β and MCP-1. As ICAM-1 and VCAM-1 are major adhesion molecules expressed on activated endothelial cells and mediated inflammatory cell infiltration into tissues, N protein may play a key role in the development of ARDS and multi-organ injury.

The N protein is highly abundant in the viruses. Its function involves entering the host cells, binding to the viral RNA genome and forms the ribonucleoprotein core to facilitate its replication and process the virus particle assembly and release^27^. Previous reports showed that the N protein from SARS-CoV and MERS-CoV were highly inflammatory nature to promote the expression of inflammatory cytokines, chemokines, prothrombinase and were able to induce acute lung inflammation in mouse model^28-30^. The N protein from hantavirus Andes virus increased basal endothelial cell permeability by activating RhoA signaling^31^. The effect of N protein from SARS-CoV-2 on host cells is less studied. In this manuscript, we report for the first time that the N protein from SARS-CoV-2 acts as a pathogen-associated molecular pattern (PAMP) to direct bind to TLR2 and activate NF-κB and MAPK signaling. In an unbiased survey of phosphorylation landscape of SARS-CoV-2 infection, SARS-CoV-2 infection is truly promoting activation of CK2 and p38 MAPK^32^. Moreover, a recent human study showed that the serum levels of soluble ICAM-1 and VACM-1 were elevated in mild COVID-19 patients, dramatically elevated in severe cases, and decreased in the convalescence phase^33^. Taken together, the current study identified N protein as a potent factor to induce endothelial activation and provided the insights to understand the phenomenon of broad endothelial dysfunction and multi-organ injury that commonly appeared in severe COVID-19 patients.

Our study suggests that targeting on N protein may benefit to prevent or treat the pathogenesis and multi-organ injury in COVID-19 patients. By screening a natural microbial compound library containing 154 natural compounds, we identified simvastatin, an FDA-approved lipid-lowering drug, as a potent inhibitor of N protein-induced endothelial activation. Several groups have raised the idea that statins can be used as early therapy to mitigate COVID-19-associated ARDS and cytokine storm syndrome^34-36^. Several recent reports showed that in-hospital use of statins is associated with a reduced risk of mortality among individuals with COVID-19^37-39^. There were more than 20 statins available in clinical use. We tested five different statins. Only simvastatin showed potent inhibitory activity on N protein-induced endothelial activation. Lovastatin also showed mild inhibitory effect, which may be due to its structure is similar to simvastatin (**Fig.6**). As there are many reports showed that simvastatin has anti-inflammatory role^40^, it would be more effective in the treatment of COVID-19 patients by multiple mechanisms. Our results justify that simvastatin may be more benefit for the treatment of COVID-19 by potently suppressing endothelial activation.

**Figure 6.**
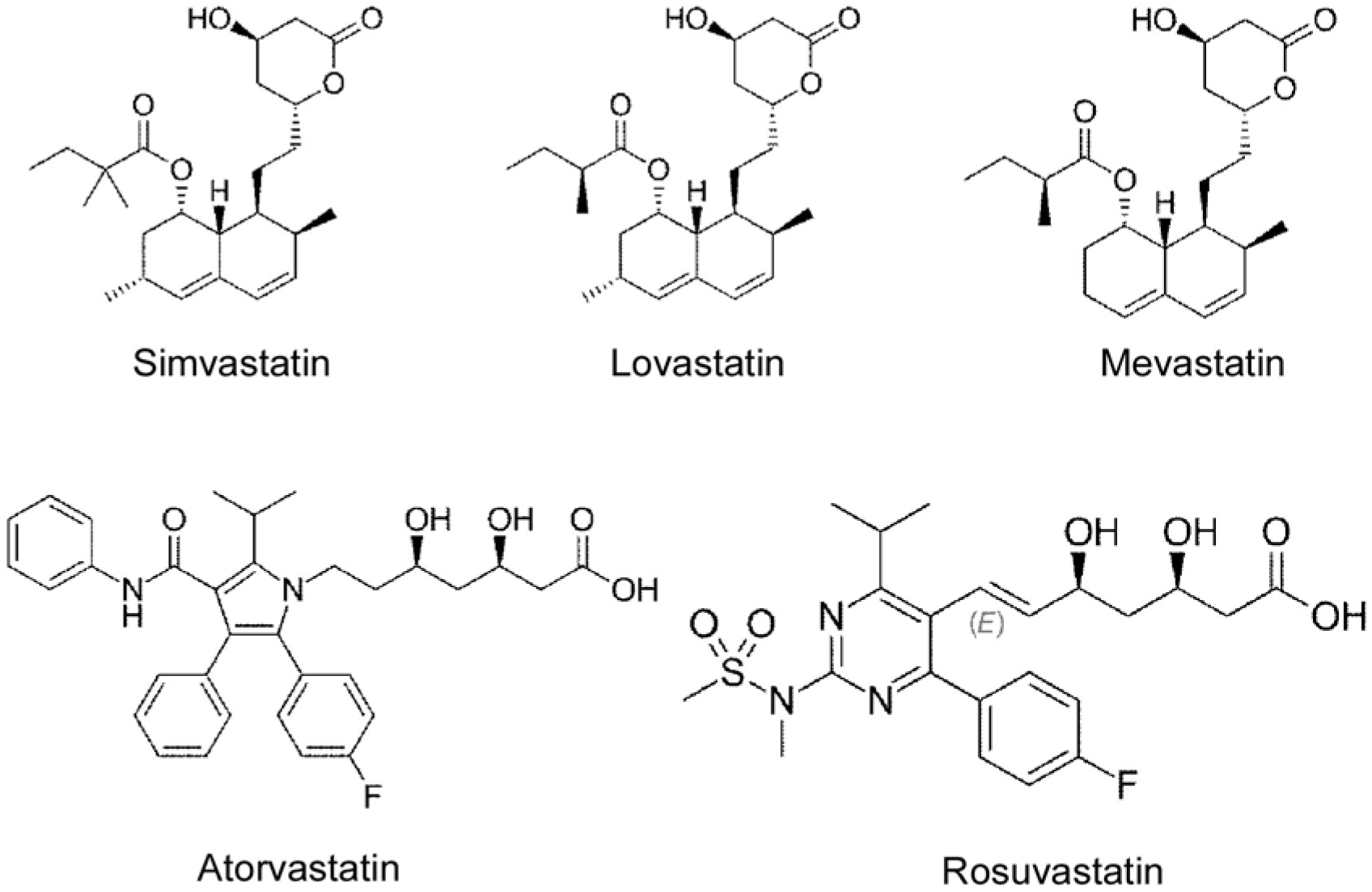
Chemical structures of Simvastatin, Lovastatin, Mevastatin, Atovastatin and Rosuvastatin (adapted from Wikipedia).

**Figure 7.**
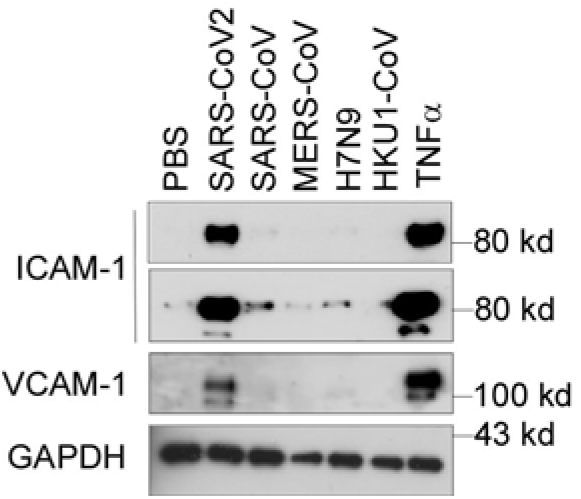
The N protein from SARS-CoV2 but not the other coronaviruses potently induced endothelial cell activation. HLMECs were treated with or without five different recombinant viral N proteins (1 μg/mL) including SARS-CoV2, SARS-CoV, MERS-CoV, H7N9 and HKU1-CoV for 8 hrs. The expression of ICAM-1 and VCAM-1 was detected by western blot. TNFα (10 ng/mL) was served as a positive control. GAPDH was served as loading control. The experiments were repeated at least one more time.

Many patients with severe COVID-19 show signs of a cytokine storm^41^. The high levels of cytokines amplify the destructive process by leading to EC activation, DIC, inflammation and vasodilation of the pulmonary capillary bed. This results in alveolar dysfunction, ARDS with hypoxic respiratory failure and ultimately multi-organ failure and death. EC dysfunction and activation likely co-determine this uncontrolled immune response. This is because ECs promote inflammation by expressing leukocyte adhesion molecules, thereby facilitating the accumulation and extravasation of leukocytes, including neutrophils and monocytes/macrophages, which enhance tissue damage. One recent report showed that SARS-CoV-2 N protein robustly induced proinflammatory cytokines/chemokines in human primary PMBCs^42^, suggesting that circulating N protein may also contribute to the initiation and progression of cytokine storm. Remarkably, though the protein sequences of N proteins from coronaviruses are highly conserved, only NP from SARS-CoV-2 induced endothelial activation. The NPs from other coronaviruses such as SARS-CoV, MERS-CoV, HKU1-CoV and influenza virus H1N1 did not affect endothelial activation. Thus, these findings are well consistent with the results from clinical investigations.

In summary, our present study identified SARS-CoV-2 N protein as a potent inducer of human endothelial activation, which can be specifically inhibited by simvastatin. The study provides insights on SARS-CoV-2-induced vasculopathy and coagulopathy, and suggests that simvastatin, an FDA-approved lipid-lowering drug, may benefit to prevent vascular pathogenesis and improve the outcome of COVID-19 patients.

## METHODS

### Reagents

SARS-CoV-2 nucleocapsid protein (NUN-C5227), S protein (SPN-C52H4) and envelope protein (ENN-C5128) were from Acrobiosystems (Newark, DE). SARS-CoV-2 NSP1 (97-095), NSP5 (10-116), NSP7 (97-096) and NSP8 (97-097) proteins were obtained from Prosci (Poway, CA). SARS-CoV-2 papain-like protease (DB604) was purchased from Lifesensors (Malvern, PA). HCoV-HKU1 coronavirus nucleocapsid protein, H7N9 Nucleocapsid Protein (40110-V08B), MERS-CoV Nucleoprotein protein (40068-V08B) and SARS Coronavirus Nucleocapsid Protein (40143-V08B) were from Sino biological (Beijing, China). Mammalian expression plasmid for SARS-CoV-2 nucleocapsid protein (152536) was obtained from addgene (Watertown, MA). ICAM-1 (60299-1-Ig) and GAPDH (60004-1-Ig) antibodies were from Proteintech (Rosemont, IL). VCAM-1 (sc-8304) and β-actin (sc-47778) antibodies were obtained from Santa Cruz Biotechnology, Inc (Dallas, TX). SARS-CoV/SARS-CoV-2 Nucleocapsid Antibody (40143-MM05) and SARS-CoV-2 Nucleocapsid antibody (40588-T62) were from Sino biological (Beijing, China). The NF-κB and MAPK signal pathway antibodies were all from Cell Signaling Technology (Danvers, MA).

### Cells

Human umbilical vein endothelial cells (HUVEC), Human Aortic Endothelial Cells (HAEC), Human Coronary Artery Endothelial Cells (HCAEC), Human Dermal Microvascular Endothelial Cells (HDMEC) and Human Lung Microvascular Endothelial Cells (HLMEC) were purchased from Lonza Bioscience (Houston, TX). Mouse lung microvascular endothelial cells were obtained from Cell Biologics Inc (Chicago, IL). Cells were cultured in different endothelial cell growth medium in a humidified incubator with 5% CO_2_ at 37°C. Endothelial cells between passages 4 and 8 were grown as a monolayer and were used in all the experiments. HLMEC were treated with vehicle (PBS) or various SARS-CoV-2 proteins, including nucleocapsid protein (NP), S protein (SP), envelope protein (EP), NSP1, NSP3, NSP5, NSP7 and NSP8 at 1 μg/mL for 8 h. In another set of experiment, HLMECs were treated with different coronavirus nucleocapsid proteins, including SARS-CoV-2 NP, SARS-CoV NP, MERS NP, H7N9 NP and HCoV-HKU1 NP at 1 μg/mL for 8 h. Different subtypes of endothelial cells were also used to observe the response to NP stimulation. For time course assay, HLMECs were incubated with 1 μg/mL SARS-CoV-2 NP for 2 h, 4 h, 8 h, 16 h and 24 h, respectively. For dose-dependent assay, HLMECs were treated with various concentrations of SARS-CoV-2 NP ranging from 0.01-10 μg/mL. For NF-κB and MAPK signal pathway assay, HLMECs were subject to SARS-CoV-2 NP exposure for 15 min, 30 min, 45 min, 1 h and 2 h, respectively. TNFα (10 ng/mL, PeproTech, Cranbury, NJ) was used in all above experiments as a positive control of endothelial activation.

### Transfection of plasmids

Flag-NP and Flag-control vectors were transfected to HLMEC by electroporation using Nucleofector device (Lonza) and Nucleofector kits (Lonza, VPB-1002) following the manufacturer’s instruction. The whole cell lysates were harvested 48 h after electroporation and were analyzed by western blot.

### Chemical screening

To screen for inhibitors of NP-induced endothelial activation, HLMECs were pretreated by 30 µM/L of individual chemical from microbial natural product library (Target Molecule, Wellesley Hills, MA) for 1 h followed by treatment of 1 µg/ml of N protein for 8 h. The effect of chemicals on endothelial activation were measured by ICAM-1 expression. To evaluate the inhibitory effects of statins on NP-induced endothelial activation, HLMECs were pretreated with atovastatin, lovastatin, mevastatin, rosuvastatin or simvastatin (all from Biovision, Milpitas, CA) for 1 h and followed by the treatment of N protein (1 µg/ml) for 8 h. The whole cell lysates were harvested and analyzed by western blot.

### Inhibitor treatment

The antagonists or inhibitors involved in NF-κB and MAPK signal pathway were introduced to verify the action of NP. The endocytosis inhibitors Pitstop 2 (12.5 μM, Sigma) and Dynasore hydrate (12.5 μM, Sigma), the TLR4 antagonist LPS-RS (10 μg/mL, InVivoGen, San Diego, CA), the TLR1/TLR2 antagonist CU-CPT22 (20 μM, Millipore, Burlington, MA), the IL-1R antagonist (20 μM, Cayman Chemical Company), IKK-16 (20 μM, Cayman Chemical Company), the JNK inhibitor V (20 μM, Cayman Chemical Company), the ERK1/2 inhibitor U0126 (20 μM, Cell Signal Technology), and the p38 inhibitor SB203580 (20 μM, Enzo Life Sciences, Farmingdale, NY) were added into HLMEC cultures 1 h before NP exposure. The whole cell lysates were harvested 8 h after NP stimulation and analyzed by western blot.

### NP-induced NF-κB and MAPK activation in 293T-TLR2 cells

293T overexpressed human TLR2 was use to confirm the interaction between NP and TLR2, Wild type 293T and 293T-TLR2 (InVivoGen) cells were cultured in DMEM supplemented with 10% FBS. The cells were treated with or without NP (1 μg/ mL) for 15 mins. The cells were harvested and the whole cell lysates were subjected to detection of pJNK and pP38 by western blot.

### QPCR

Total RNA was isolated from HLMEC after 8 h of NP exposure with the RNeasy Mini Kit (Qiagen, Germantown, MD) according to the manufacturer’s instructions. The first-strand cDNAs were synthesized by the High Capacity RNA-to-cDNA Kit (Thermo Fisher Scientific, Vilnius, Lithuania). The reaction mixture contained 2×SYBR Green PCR Master Mix (Applied Biosystems, Foster City, CA, USA), primer pairs and cDNAs. The reaction consisted of a 2-step thermocycling protocol (95 °C for 15 s and 60 °C for 1 min; 40 cycles). The mRNA levels were calculated by using the 2^−ΔΔCT^ method. The Primer sequences used in the experiment were listed in Table S1. Results were obtained from at least three biological replications performed in triplicate.

### Western blot

The total protein was collected with ice cold RIPA lysis buffer after NP stimulation. Equal amounts of protein were loaded into the wells of the SDS-PAGE gel and separated by electrophoresis. Then the protein were transferred to the PVDF membrane. The 5% (w/v) skim milk were used to block the un-specific binding. Membranes were incubated with different primary antibodies at 4 °C overnight, followed by incubation with HRP-conjugated secondary antibodies for 1 h at room temperature. Luminescence was generated after the membranes were exposed to Super Signal West Pico Chemiluminescent Substrate (Thermo Fisher Scientific) and was detected with X-ray film.

### Monocyte adhesion assay

Monocyte adhesion was analyzed as previously described^43^, with some modification. HLMECs were stimulated with NP or TNF-α for 8?h. The human acute monocytic leukemia cell line THP-1 (Lonza) was prelabeled with Zombie Red fluorescent dye (Biolegend, San Diego, CA) in RPMI-1640 medium for 30?min at 37°C before being added to HLMECs and co-cultured for 1 h. Non-adherent cells were removed by gently washing with cold RPMI-1640 medium. The images of adherent THP-1 cells were taken under Cytation 3 Cell Imaging Multi-mode Reader.

### Statistical Analyses

Data were obtained from at least three independent experiments and were represented as mean ± SD. Statistical differences were compared using one-way ANOVA followed by a Tukey post hoc test. An unpaired Student’s t-test was performed to compare data between two independent groups. A difference with P < 0.05 was considered statistically significant. A p-value less than 0.05 was considered statistically significant.

## ACKNOWLEDGMENTS

We thank Dr. Peter Klein (University of Pennsylvania) for providing SARS-CoV2 Nucleocapsid plasmid via Addgene.com. This work was supported by National Institutes of Health Grant AI138116 (to M.F). J.Q. was supported by National Institute of Health Grant AI150877 and AI144564. Y.Q was supported by China Scholarship Council (20190682503)

## AUTHOR CONTRIBUTIONS

M.F. and Y. Q designed the research and analyzed data; Y.Q., T.L., and P.P did the experiments; C.L, P.N., J.Q and H-B. X provided advice and critically read the manuscript; M.F and Y.Q wrote the manuscript.

## DECLEARE OF CONFLICT OF INTERESTS

The authors have no conflict of interests to declare.

---

**Table S1.**
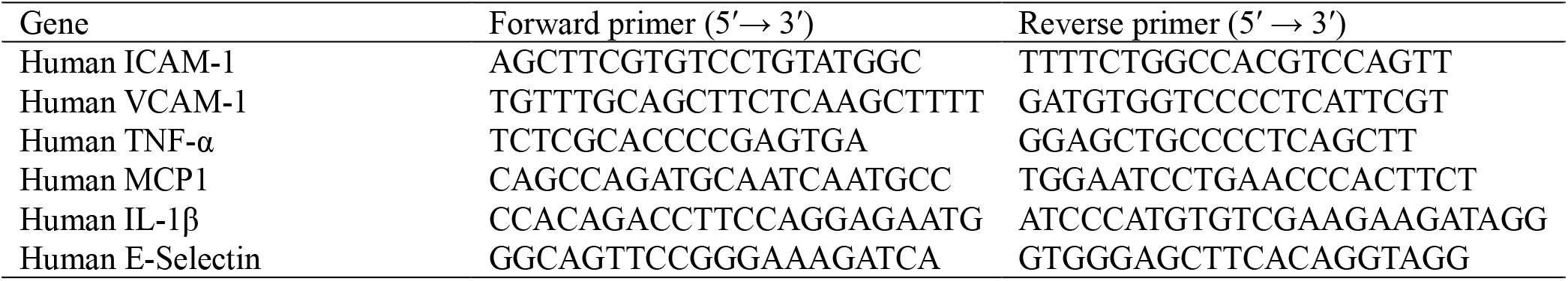
Primers used in QPCR reactions

